# Dynamic tracing using ultra-bright labelling and multi-photon microscopy identifies endothelial uptake of poloxamer 188 coated poly(lactic-co-glycolic acid) nano-carriers *in vivo*

**DOI:** 10.1101/2020.11.19.385062

**Authors:** Igor Khalin, Caterina Severi, Doriane Heimburger, Antonia Wehn, Farida Hellal, Andreas Reisch, Andrey S. Klymchenko, Nikolaus Plesnila

## Abstract

Poly(lactic-co-glycolic acid) (PLGA)-based drug formulations are approved for the use in humans, however, the potential of PLGA to design nanoparticles (NPs) and target the central nervous system remains to be exploited.

The aim of the current study was design PLGA NPs, loading them with bulky fluorophores thereby increasing single particle fluorescence to a level visible by *in vivo* microscopy, and investigate their brain biodistribution. We developed, highly fluorescent 70 nm PLGA NPs significantly brighter then quantum dots enabling their visualization by intravital real-time 2-photon microscopy. We found that PLGA NPs coated with pluronic F-68 (PF-68) had a substantially longer plasma half-life than uncoated NPs and were taken up by cerebro-vascular endothelial cells. High resolution confocal microscopy revealed that coated PLGA NPs were present in late endothelial endosomes of cerebral vessels within 1 hour after systemic injection and were more readily taken up by endothelial cells in peripheral organs.

The current data suggest that PF-68 coated PLGA NPs are taken up by mouse cerebral and peripheral endothelial cells *in vivo*. The combination of ultra-bright NPs and *in vivo* imaging may thus represent a promising approach to reduce the gap between development and clinical application of nanoparticle-based drug carriers.

## 1. Introduction

Polymeric nanoparticles (NPs) are promising carriers for the targeted delivery of drugs and may thus significantly improve pharmacotherapy. Among these, poly(lactic-co-glycolic acid) (PLGA) NPs are of specific interest since *in vivo* PLGA is hydrolyzed into the physiologic metabolites lactic and glycolic acid and is therefore regarded to have a low toxicity and to be biocompatible and biodegradable [1]. Due to these properties, PLGA-based injectable formulations have been approved for human use by the United States Food and Drug Administration and by the European Medicine Agency [2]. Their market is expected to reach about 930 billion in 2024 [3] and the FDA has launched a dedicated Regulatory Science Program directed to improve bioequivalence of PLGA-based drug products [4].

Despite these significant advances, it is still particularly difficult to target drugs to the brain [5]. Nano-formulations may overcome these difficulties, but despite substantial efforts, the clinical translation of NPs able to specifically deliver pharmacologically active substances across the blood-brain barrier is still very poor [6]. For example, it was demonstrated that poloxamer 188 (Pluronic F-68, PF-68) coated PLGA nanoparticles may improve CNS drug delivery [7–11], however, the exact mechanism of nanoparticle-mediated uptake of drugs into the brain remains elusive [12].

One of the reasons responsible for this situation is the limited ability to directly track individual NPs *in vivo* [11]. NPs-based drug carriers are too small to be observed by light microscopy. Therefore, the main strategy employed to overcome this issue was to make fluorescent NPs so bright, that their signal became detectable. Several approaches were developed to increase the brightness of polymer based fluorescent nanoparticles, either using conjugated polymers, like in p-dots [13], or by encapsulating large amounts of dyes [14]. In the latter case, however, high dye concentrations caused self-quenching and reduced particle fluorescence. This issue was finally solved by aggregation-induced emission dyes [15] or, as developed by our team, co-loading NPs with bulky hydrophobic counter-ions [16]. The very high brightness of such NPs allowed us to image individual NPs in cultured cells down to the subcellular level [17–19] and to use Förster Resonance Energy Transfer (FRET) for detection of biomolecules at the single molecule level [20–22].

In the current animal study, we aimed to apply the technology of excitation energy transfer (EET) between bulky dyes to generate high fluorescence in NPs with a clinical potential, *i.e.* PLGA NPs. Using PLGA NPs bright enough to have the potential to become visible *in vivo*, we aimed to investigate the properties of uncoated or PF-68 coated PLGA NPs in suspension, on surfaces, and *in vivo*. To do so, we first optimized the brightness of PF-68 coated PLGA (PF68-PLGA) NPs by varying the type of dyes, dye loading strategy, and particle size. Then we investigated the biodistribution of these optimized PLGA NPs in the mouse brain *in vivo* by 2-photon microscopy (2PM). Subsequently, we correlated the findings obtained *in vivo* with data from fixed tissue obtained by high-resolution confocal microscopy (CM) and 3D-reconstruction to unambiguously determine the spatial distribution of individual NPs *in vivo*.

## 2. Results

### 2.1 Formulation and characterization of nanoparticles

Dye-loaded PLGA NPs were assembled through nanoprecipitation [23, 24]. A solution of the polymer and the dye in a water-miscible solvent (here acetonitrile) was quickly added to a large excess of an aqueous phase, resulting in supersaturation and formation of nanoparticles **(Figure 1A)**. We chose two types of dye systems with similar spectra for encapsulation: the dye salt R18/F5-TPB, known to be very bright, due to counter-ions which prevent aggregation and self-quenching [16, 25] and the bulky perylene dye lumogen red, which combines very high brightness and photostability [26]. Based on previous data, we chose 1 wt% of the dyes with respect to the polymer as optimum loading. The most straightforward approach to increase NPs brightness is to increase their size, since the number of encapsulated dyes increases with the third power of the radius. A tenfold increase in brightness can thus be achieved by a 2.2-fold increase in size. In the case of nanoprecipitation the particle size depends on the concentration of the polymer, but also on the interactions of charges [27]. Here, we used a polymer concentration of 4 g·L^−1^ and a buffer solution containing NaCl to increase particle size [23, 28]. The role of the NaCl is to screen interactions between charges on the polymer, which influences particle formation, and leads to larger particles [18, 28]. The used combination allowed increasing the particle size from about 30 nm under reference conditions (2 g·L^−1^, 20 mM phosphate buffer, pH 7.4, no additional salt) to around 70 nm, while maintaining a low polydispersity index (PDI) ≤ 0.1 **(Table S1)**. Interestingly, R18/F5-TPB led to slightly bigger particles, which could be further tuned by adjusting the salt concentration. Thereby, we were able to produce PLGA NPs loaded with lumogen red or R18/F5-TPB with diameters of 70 to 80 nm **(Figure 1B)**.

**Figure 1.**
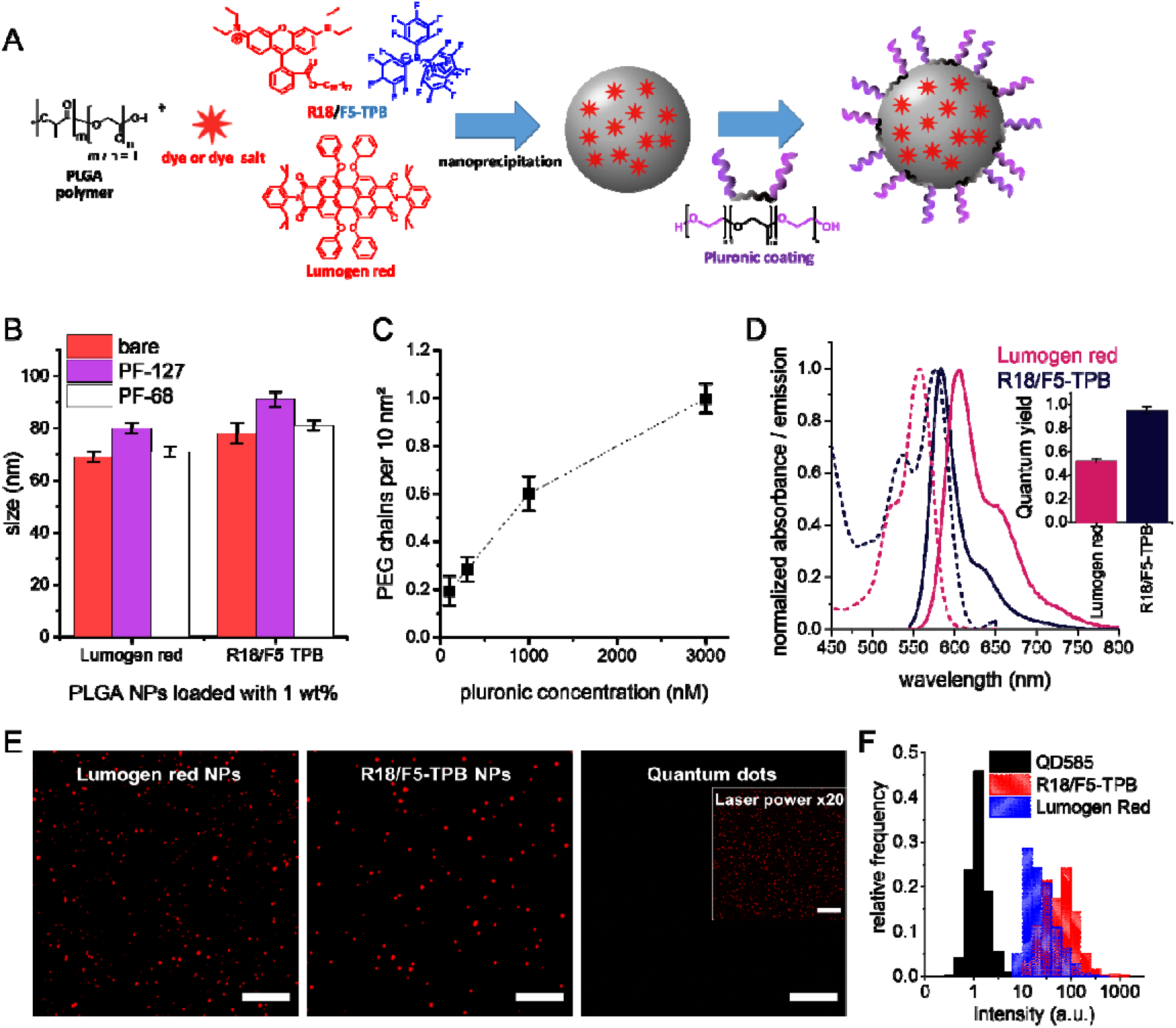
Formulation and analysis of bright PLGA nanoparticles. **A:** Schematic drawing of the assembly of biodegradable dye-loaded PLGA NPs through nanoprecipitation followed by coating with Pluronics. **B:** Sizes of NPs loaded with 1 wt% lumogen red or R18/F5-TPB, either bare or coated with 3000 nM of the Pluronics F-127 or F-68, as obtained by DLS. **C:** Density of PEG chains on the NPs surface at different pluronic concentrations as obtained from FCS measurements using lissamine labelled Pluronic F-127 after dialysis for 24h. Error bars correspond to standard error of the mean for at least three independent measurements. **D:** Normalized absorption (dashed lines) and emission (solid lines) spectra of PLGA NPs. Error bars correspond to three independent measurements. **E:** Fluorescence micrographs of NPs adsorbed on a glass surface. Excitation was 550 nm. To make quantum dots (QDs) visible, the laser power had to be increased by 20 times (insert). Scale bars correspond to 10 μm. **F:** Histograms of particle intensity distribution obtained from based on images from panel (E).

In order to stabilize the particles in biological media and to reduce non-specific interactions, we then adsorbed two types of pluronics, F-127 and F-68, onto the particles. PEGylated particles have been shown to have increased stability in salt and buffer solutions and strongly reduced protein adsorption [18, 27] leading to prolonged circulation half-lives *in vivo* [29]. We investigated the adsorption of Pluronics with fluorescence correlation spectroscopy (FCS) using Pluronic F-127 (PF-127) labelled with lissamine. This allowed us to directly measure the number of adsorbed pluronics per single NP (here NPs of 30 nm were used in order to avoid perturbations of the FCS measurements) depending on the added concentration **(Figure 1C)**. The density of pluronics and, in consequence, the density of PEG chains on the surface of NPs increased continuously up to a concentration of 3000 nM. At this concentration about 100 pluronics per NP remained on the surface after dialysis. This corresponds to about one PEG chain per 10 nm^2^, a value well in line with previous results [30]. Adsorption of pluronic PF-127 or PF-68 onto the surface of NPs increased the particle size by 10 and 4 nm, respectively **(Figure 1B)**. This increase in size indicates the formation of a PEG shell around the NPs.

Encapsulation of the dyes within NPs resulted in absorption and emission spectra similar to those of the dyes in solution, indicating minimal aggregation of the dyes within NPs **(Figure 1D, Figure S1)**. The quantum yields (QY) of R18/F5-TPB and lumogen red remained at a very high level (95 and 52%, respectively) as determined from the absorbance and fluorescence spectra, using Rhodamine 101 as reference [31]. Based on the particle size and loading, it can be estimated that particles contained about 1000 dyes in the case of lumogen red and 1150 in the case of R18/F5-TPB. Using the absorption coefficients of the dyes (ε ≈ 100,000 M^−1^.cm^−1^ for R18/F5-TPB and ≈75,000 M^−1^.cm^−1^ for lumogen red) the particle brightness can be estimated by N×ε×QY, where N is the number of dyes per particle: 1.1 ×10^8^ M^−1^.cm^−1^ for the R18/F5-TPB NPs and 4.0 × 10^7^ M^−1^.cm^−1^ for the lumogen red NPs. This brightness can be compared to that of quantum dots of similar color (here QD585 from Invitrogen), a common, commercially available standard for bright NPs. At an excitation wavelength of 550 nm (using our microscopy setup) the QD585 brightness was about ~2.1×10^5^ M^−1^cm^−1^ (estimated form the data of provider), *i.e.* at least two orders of magnitude higher than QDs. It should be noted that QD585 exhibit a ~10-fold higher absorbance (and thus higher brightness) in the violet spectrum (405 nm), but for a direct comparison with our NPs, we used the identical conditions. We further compared single-particle brightness by wide-field fluorescence microscopy, after deposition of the particles on glass surfaces at dilutions maximizing the deposition of individual particles [16]. This comparison revealed that lumogen red and R18/F5-TPB NPs have a 23 and 55 times higher brightness than quantum dots, respectively **(Figure 1E and F)**. The lower values obtained here are due to a combination of multiple factors. The first one is an overestimation of the particle size (and thus the number of dyes per particle) by DLS, which assesses hydrodynamic diameter rather than size [22] of the particle core. The second one is a relatively strong irradiation power used during microscopy (as compared to a fluorimeter), which may lead to saturation of the multi-chromophore system due to singlet-singlet annihilation [32].

### 2.2 Intravital evaluation of systemic circulation of PLGA NPs

Our next step was to investigate whether lumogen red loaded PLGA NPs coated with PF-68 can be visualized *in vivo*. We implanted an acute cranial glass window over the cerebral cortex of a mouse, inserted a femoral arterial catheter for systemic injection of NPs, and positioned the animals under a 2-photon microscope as previously described [29]. Such a system allowed us to inject the NPs solution into the mouse directly under the microscope. After visualizing cerebral vessels with a plasma tracer (fluorescein isothiocyanate (FITC)-dextran 2000 kDa; **Figure 2A; Baseline**), we chose a region of interest (ROI) and injected PLGA NPs loaded with lumogen red with (PF68-PLGA) and without PF-68 coating (PLGA bare). In both groups we immediately observed circulating PLGA NPs in the brain vasculature **(Figure 2A; 0 min)**. Sixty minutes after injection, bare NPs were almost undetectable, while coated NPs were clearly visible **(Figure 2A; 60 min)**. Despite the fact that bare and coated NPs were injected at equal doses (7.5 μL·g^−1^), the absolute fluorescence intensity of coated NPs was significantly higher than the one of bare NPs right after injection **(Figure 2B)**. Quantification of relative intravascular fluorescence intensities over time showed that intensity of bare NPs dropped steeply to 35% of baseline already 5 minutes after injection and became negligible 25 minutes later. In contrast, 63% of the initial fluorescence intensity of PF68-PLGA NPs was still present even one hour after injection **(Figure 2C)**. These *in vivo* data demonstrate that lumogen red-loaded PLGA NPs coated with PF-68 are sufficiently bright and have a blood half-life for more than 1 h.

**Figure 2.**
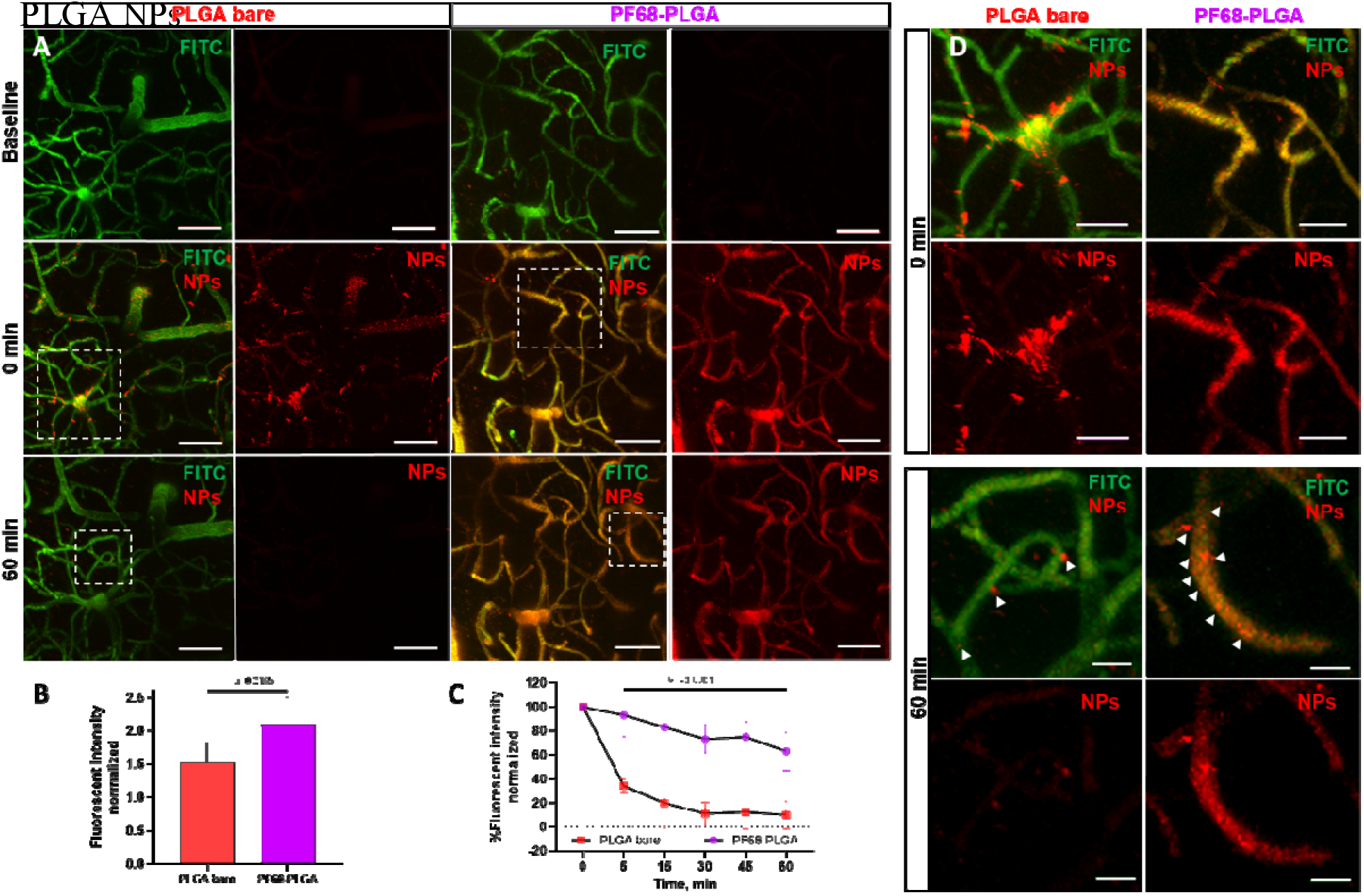
Intravital 2-photon microscopy of PLGA NPs in mouse brain vasculature. **A:** 2-photon microscopy (2PM) images after injection of FITC-dextran 2000kDa to label blood plasma (Baseline) and immediately (0 min) or 60 minutes after additional injection of bare or PF-68 coated PLGA (PF68-PLGA) NPs. Scale bars: 20 μm. **B:** Quantification of fluorescent intensity (FI) of circulating coated and not coated PLGA NPs immediately after injection. FI was normalized to FITC and baseline. **C:** Temporal kinetics of the FI of bare vs. PF68-PLGA NPs quantified from 2-PM images. Data presented as means ± standard deviation. Statistical analysis: two-way ANOVA with Tukey correction for multiple comparison (bare NPs n=3, coated NPs n=6). **D:** High power images of the regions indicated by the dotted squares in A. White arrows: NPs stuck in the vessel wall. Scale bars: 10 μm.

Analysis of higher magnification images revealed that PLGA bare NPs presented as dispersed heterogeneous aggregates with varying fluorescence intensities; some aggregates seemed to stick to circulating blood cells **(Figure 2D, left; 0 min and Movie S1)**. In contrast, PF68-PLGA NPs produced a homogenous fluorescence signal within the vascular lumen and circulated without any obvious stalls early after injection **(Figure 2D, right; 0 min and Movie S2)**. Due to the high velocity of cerebral blood flow and the low maximal frequency of the 2-PM scanner, coated particles could not be detected individually. One hour after injection, practically no bare NPs could be observed **(Figure 2D, left; 60 min)**, while coated PLGA NPs were still present within the vessel lumen and emitted a bright fluorescence **(Figure 2D, left; 60 min)**. Interestingly, we detected that few uncoated and a large number of coated PLGA NPs did not circulate, but became associated with the vessel wall, suggesting adhesion to the endothelial surface or endothelial uptake **(Figure 2D, 60 min, white arrows)**. We followed the NPs for another 60 min, *i.e.* until 120 minutes after injection, but did not observe major changes **(Figure S2)**; PLGA NPs were still aligned with the vessel wall.

### 2.3 Analysis of the PF-68 PLGA NPs bio-distribution in post-fixed brain

Since 2-PM microscopy does not have the necessary resolution to distinguish between adhesion of NP to the vessel wall and uptake of NPs by endothelial cells, we continued our investigations using perfusion-fixed brain tissue and high-resolution confocal microscopy. For this purpose, naïve mice, *i.e.* mice without a cranial window, were sacrificed 60 minutes after intravenous injection of PBS (control), bare, or coated PLGA NPs **(Figure S3A)**. Then, coronal brain sections **(Figure S3B-D)** were evaluated histologically for particle fluorescence. In whole brain tiles and plane scans (**Figure 3A and B**) we detected that the red fluorescent signal was more intense in animals which received PLGA NPs. Moreover, the intensity in the brain was 2-fold higher for coated NPs compared with PBS (**Figure 3C**).

**Figure 3.**
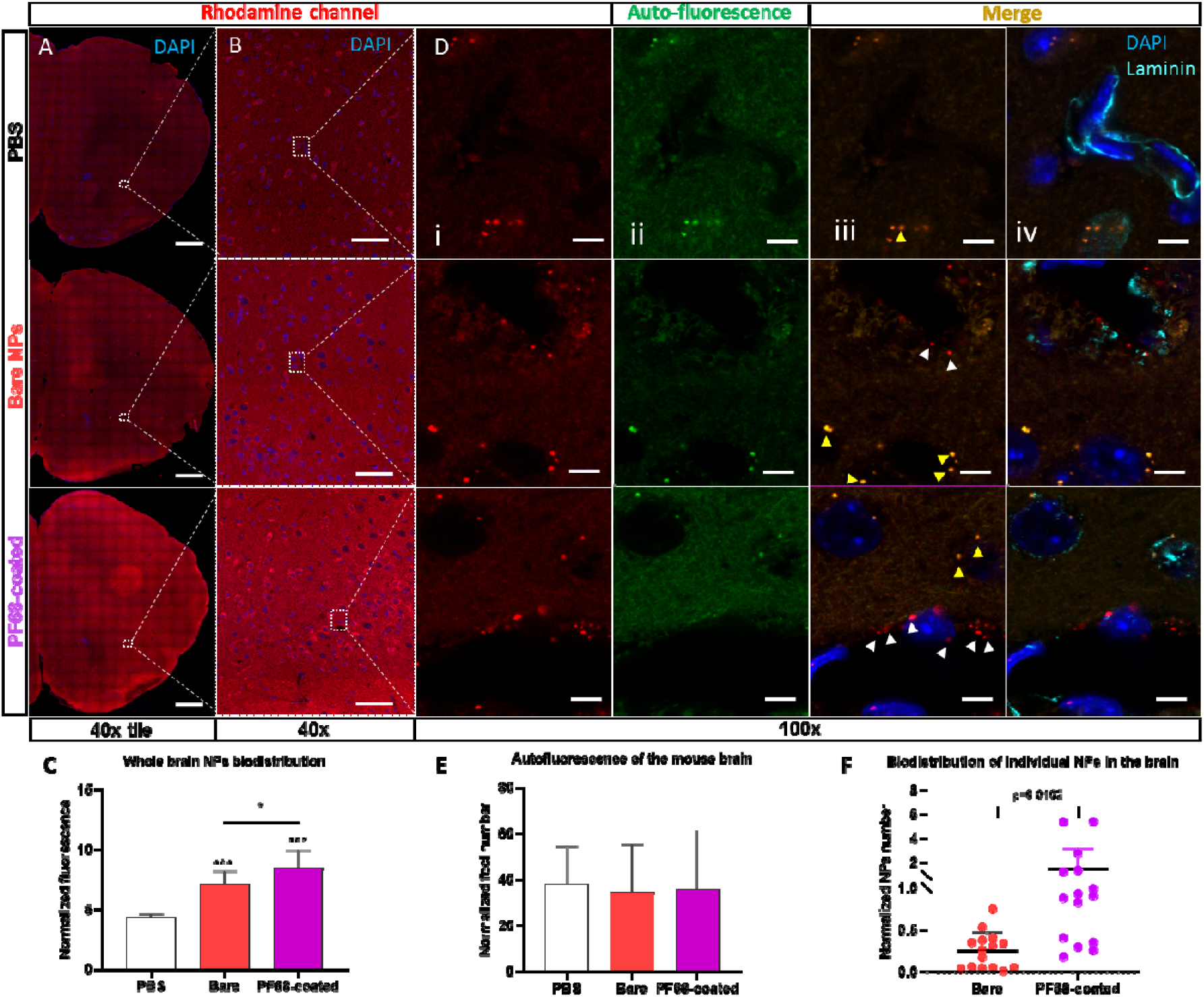
Biodistribution of F68-coated NPs in the mouse brain 60 minutes after systemic application. Confocal microscopy images of coronal brain section recorded with an 40x objective from mice which received PBS, bare, or coated red fluorescent PLGA NPs. **A:** Tile scan of whole section and **B:** a single scan. Nuclei were stained with DAPI. Scale bars: 1mm (40x tile), 50 μm (40x single). **C:** Quantification of the distribution of the NPs throughout the whole brain, analyzed using FIJI: the mean grey value of the rhodamine channel normalized to the mean grey value of the DAPI channel. Data presented as means ± standard deviation (SD). For the statistical analysis, one-way ANOVA was used followed by multiple comparison with the Tukey correction. *** p◻0.001 compared to PBS group; * p◻0.05 compared to F68-coated group; (n=3 animals x 3 whole brain regions (rostral, middle, caudal)). **D:** A zoomed single plane from **B** of brain parenchyma, stained by Laminin and DAPI. **i:** Rhodamine channel shows non-specific red foci. **ii:** Auto-fluorescent channel shows auto-fluorescent foci of the brain parenchyma. **iii**: Merge channels show true NPs signal which appeared only in rhodamine channel. White arrows indicate the PLGA NPs, while yellow arrows indicate auto-fluorescence as a co-localization between rhodamine and auto-fluorescent channels. **iv:** Basal lamina stained by Laminin outlines the spatial distribution of NPs. Scale bar - 5 μm. **E:** Quantification of the autofluorescent foci (co-localized with autofluorescent channel) and **F:** PLGA NPs distribution (non-colocalized with autofluorescent channel). Foci or PLGA NPs found in the region of interest (ROI) was normalized to the ROI area (μm^2^) and cell numbers (n=3 x ROI=5). Data presented as mean ± SD. For the statistical analysis of autofluorescence, one-way ANOVA was used followed by multiple comparison with the Tukey correction; for the NPs - unpaired t-test.

Images taken at higher magnification (100x) showed discrete red fluorescent foci not only in tissue from mice which received PLGA NPs, but also in control animals (**Figure 3D, i**). When switching to the green channel (**Figure 3D, ii**), it became apparent that some of the fluorescent foci in the red channel also emitted green fluorescence, indicating auto-fluorescence. In fact, most of the auto-fluorescent signals were located around nuclear structures. When merging the red and the autofluorescence (green) channels (**Figure 3D, iii**), all fluorescent signals observed in brain tissue from control mice turned out to colocalize and have thus been identified to be caused by auto-fluorescence (**Figure 3D, iii, yellow arrows**), while in animals which received PLGA NPs also red, (non-colocalized) foci without a corresponding signal in the green channel were observed (**Figure 3D, iii, white arrows**). Using this approach, we were able to unambiguously identify individual PLGA NPs in brain tissue.

To further unravel the cellular distribution of PLGA coated NPs in the brain, we stained the basal lamina with anti-laminin antibodies (**Figure 3D, iv)**. After 3D-reconstruction, we observed that bare and coated PLGA NPs were located inside the luminal part of the basement membrane **(Movie S3)**, suggesting that NPs were not able to cross the BBB, but were internalized by endothelial cells. Taken together, using high-resolution confocal microscopy, correction for unspecific auto-fluorescence, and 3D-reconstructions, we demonstrate that PF68-PLGA NPs accumulate within the brain by being entrapped in endothelial cells.

Furthermore, quantitative analysis of the brain sections described above, demonstrated that mice, which received red fluorescent PLGA NPs, emitted significantly more fluorescence than control brain tissue (**Figure 3C**), suggesting accumulation of NPs in the brain. In addition, quantification of auto-fluorescence in the same tissue sections, showed that, as expected, all experimental groups emitted the same amount of auto-fluorescence, *i.e.* fluorescence not deriving from NPs (**Figure 3E**). When correcting the total particle count for the auto-fluorescence signal, it became apparent that PF68-coated NPs accumulated at 8-times higher numbers than uncoated NPs (**Figure 3F**). As such, this method could improve the signal-to-noise ratio and clearly detect that coating with PF-68 may facilitate the accumulation of NPs within the brain. However, we do not know whether by an enhanced specific interaction of coated NPs with the endothelial cell membrane, like commonly suggested [10–12], or merely by an increased contact time with the vessel wall due to a longer plasma half-life **(Figure 2C)**.

In order to investigate the bio-distribution of coated NPs at the subcellular level, we stained late endosomes, the cellular structures suspected to be involved in the uptake of NPs from the extracellular space, using the specific marker lysosomal-associated membrane protein 1 (LAMP-1; **Figure 4A)**. Using the same technique, to avoid unspecific signals **(Figure 4B, i)** as described above, *i.e.* adding auto-fluorescence signals to the particle-specific rhodamine channel, we were able to clearly distinguish between yellow auto-fluorescent structures **(Figure 4B, ii, yellow arrow)** and red NP-specific signals within capillary endothelial cells **(Figure 4B, ii, red arrows)**. When co-localizing NP-specific signals with LAMP-1, we identified NPs located in late endosomes **(Figure 4A, iii, magenta arrows)**. To confirm this colocalization, we took advantage of the depth information obtained by confocal microscopy and performed 3D-reconstructions of the area of particle accumulation (**Figure 4C, Movie S4**). Indeed, we found a large number of empty lysosomes (**Figure 4C, white spheres**) and lysosomes emitting autofluorescence (**Figure 4C, red/green spheres, yellow arrow**), however, next to free NPs (**Figure 4C, red spheres, red arrows**), we also found NPs co-localizing with the LAMP-1 signal (**Figure 4C, red/white spheres, magenta arrows**). All in all, these findings clearly suggest, that PLGA NPs were taken up by brain endothelial cells and ended up in late endosomal compartments.

**Figure 4.**
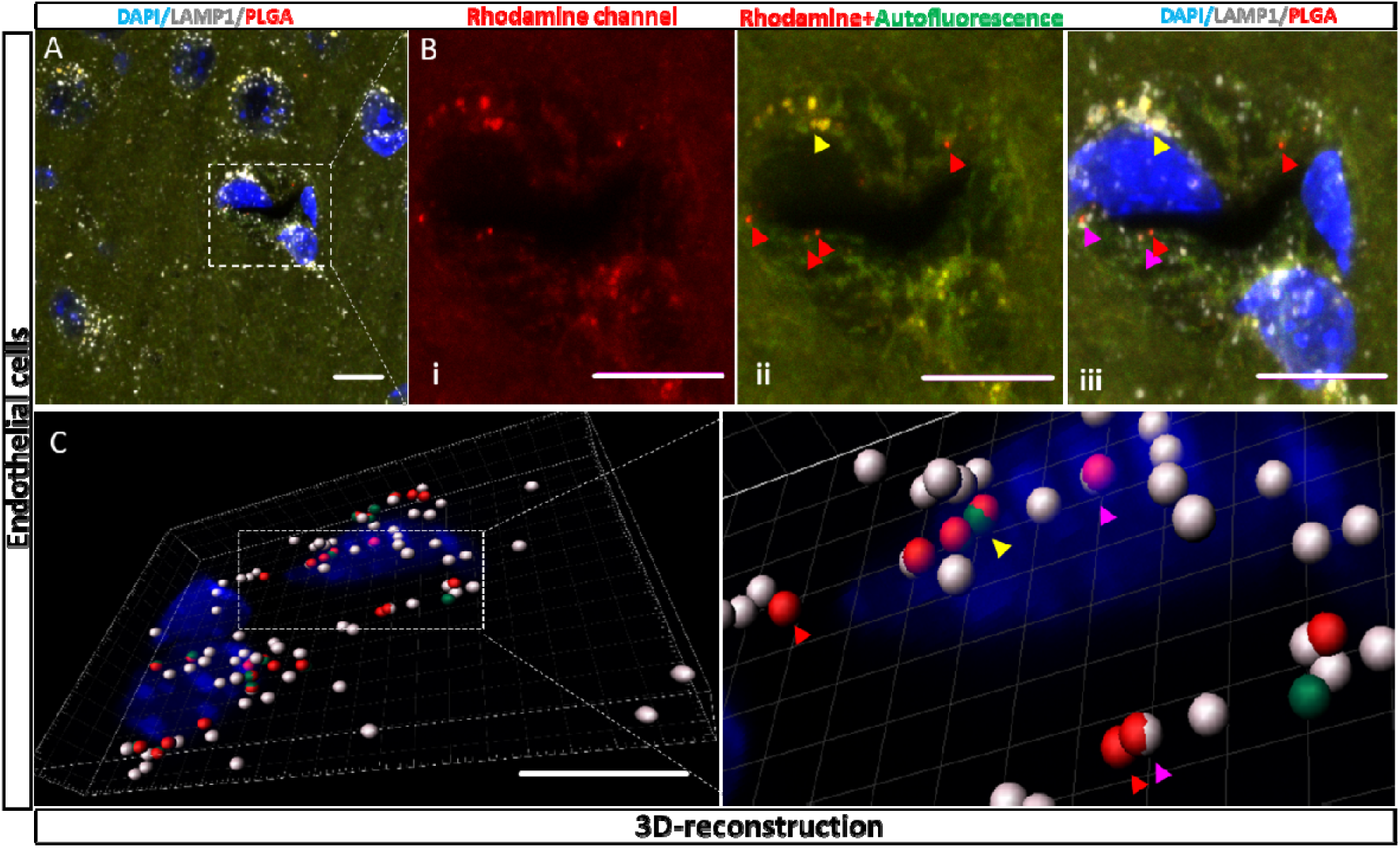
Intracellular distribution of coated PLGA NPs in brain capillary endothelial cells. **A**: A brain capillary identified by its lumen and three nuclei (blue) of vessel wall cells. Vascular cells contain a number of lysosomes as identified by LAMP-1 antibodies (white). **B**: Rhodamine channel. The vessel wall emits a large number of red particle-like fluorescence signals. **ii**: Green auto-fluorescence channel merged with rhodamine channel. Auto-fluorescence (yellow arrow) and red fluorescence emitted by NPs (red arrow). **iii:** LAPM-1 signal suggests that next to free NPs (red arrows), some NPs are taken up into lysosomes (magenta arrows). **C:** 3D reconstruction of the image shown in **A.** White spheres indicate LAMP1 positive cell compartments, red spheres NPs, and green spheres auto-fluorescent structures. Most lysosomes are empty, however, next to lysosomes emitting auto-fluorescence (red-green spheres, yellow arrow) a number of lysosomal signals also colocalise with NPs fluorescence (red-white spheres, magenta arrows). Scale bars: 5 μm.

Then, we explored whether uptake of NPs by endothelial cells was a brain specific event or if it also occurred in other tissues. For this purpose, we stained liver, spleen, kidney, and heart with collagen IV, a specific marker of the vascular basement membrane [33] **(Figure 5A)**. We found significant uptake of both bare and coated PLGA NPs in these organs **(Figure 5A)**. By quantitative analysis using the Manders colocalization coefficient [34] **(Figure 5A, B)**, we identified that the level of colocalization of coated NPs with vessels was significantly higher compared to bare NPs in all investigated organs **(Figure 5B)**.

**Figure 5.**
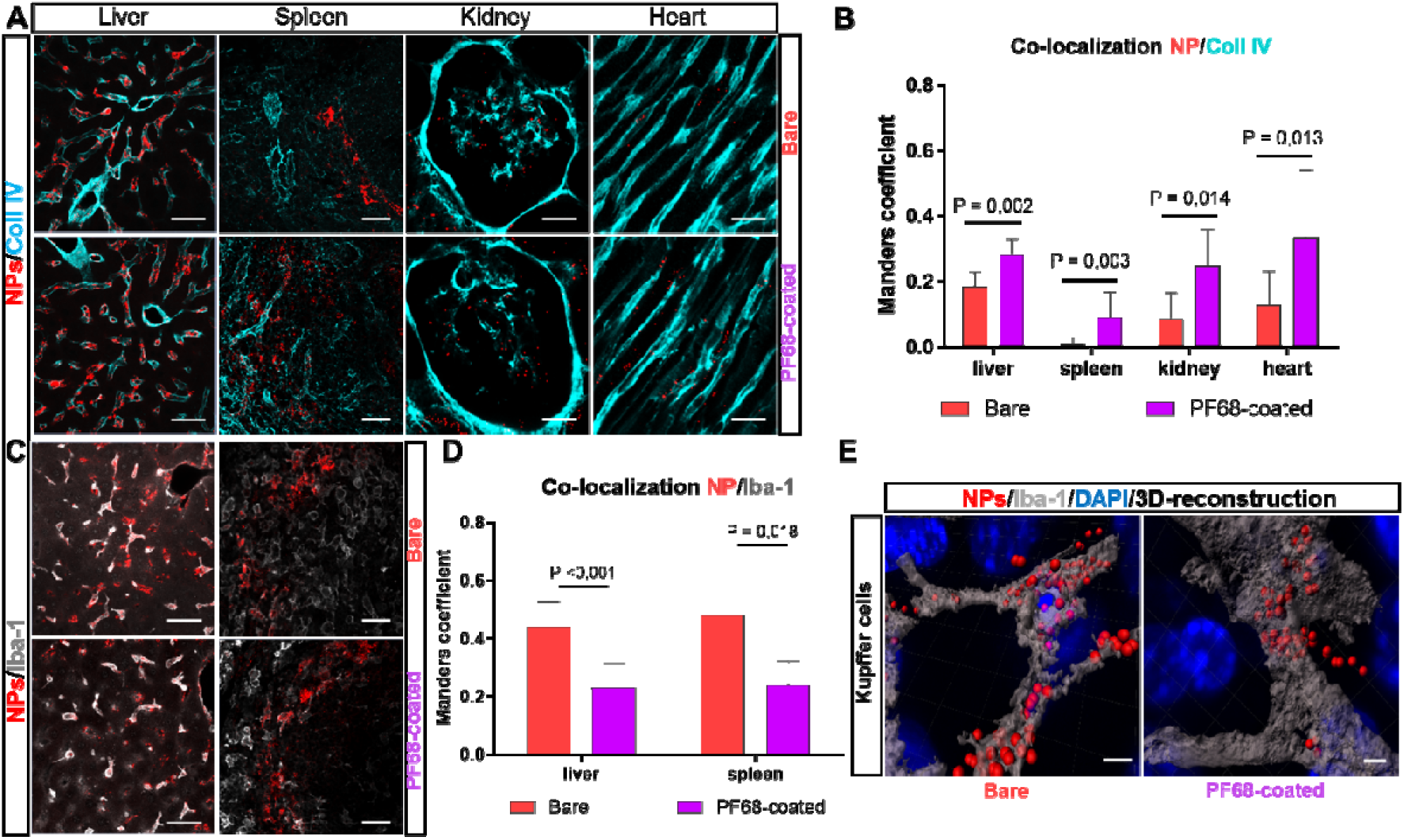
Uptake of PLGA NPs by endothelial and immune cells of the liver, spleen, kidney and heart. Representative confocal images of collagen IV **(A)** and Iba-1 **(C)** stained sections of from animals injected with bare and coated NPs. Maximum intensity projection of merged rhodamine and 647 nm channels. Scale bar: 20 μm. Quantification of co-localization of bare and coated NPs with collagen IV **(B)** and Iba-1 **(D)** using the FIJI JACoP Plugin. For the statistical analysis, Mann-Whitney Rank Sum Test was used; n=3 mice and 3-4 ROIs per section. **E:** Representative 3D reconstruction of 100x confocal images of Kupffer cells containing bare (left) and coated (right) NPs. Blue – DAPI, red spheres – PLGA NPs, grey surface – Kupffer cell. Scale bar: 3 μm.

Hence, coating promotes NPs to accumulate not only in the brain, but also in peripheral endothelial cells.

Finally, we investigated colocalization of NPs with ionized calcium-binding adapter molecule 1 (Iba-1), a specific marker for phagocytic cells [35], *i.e.* Kupffer cells in the liver and red pulp macrophages in the spleen. We observed significant colocalization of bare and coated NPs fluorescence with iba-1 phagocytic cells in liver and spleen tissue. Interestingly, bare NPs were more readily taken up by macrophages than coated ones **(Figure 5C, D)**. To evaluate this finding on a cellular and subcellular level, we performed 3D reconstructions of individual Kupffer cells. This high-resolution approach unambiguously demonstrated a higher number of bare PLGA NPs inside the cytoplasm of Kupffer cells; the uptake of PF-68 coated NP was much less efficient **(Figure 5E, Movie S5, S6)**, suggesting that PF-68 coating decreased phagocytosis of NPs on the level of each individual macrophage.

Taken together, *in vivo* imaging of the brain by 2-photon microscopy and high-resolution immunohistochemistry demonstrate that NPs are compartmentalized in endothelial late endosomes and neither bare nor coated PLGA NPs reach the brain parenchyma. Further, our results demonstrate that outside the brain bare PLGA NPs are readily taken up by phagocytic cells, *e.g.* in liver and spleen. Coating with PF-68 increases endothelial uptake and reduces phagocytosis by macrophages significantly.

## 3. Discussion

We significantly increased the fluorescent intensity of PLGA NPs by loading bulky fluorescent dyes: lumogen red or rhodamine B derivative with bulky counter-ions. Thereby, we obtained PLGA NPs with a fluorescence 23-55 times higher than quantum dots (QD585 at 550 nm excitation), the current golden standard, and were able to directly visualize their spatial-temporal biodistribution *in vivo* by 2-photon and confocal microscopy. Using this approach, we demonstrate that, within 1 hour, PF-68-coated PLGA NPs are compartmentalized in late endosomes of cerebro-vascular endothelial cells, *i.e.* NPs do not enter the brain parenchyma. Additionally, bare PLGA NPs are readily taken up by macrophages in liver and spleen and coating with PF-68 shifts their uptake towards endothelial cells.

When evaluating the pharmacokinetics of fluorescently labelled NPs *in vivo*, false positive results are a common issue due to leaking of fluorophores from NPs and unspecific binding to other structures. For example, up to 28-68% of Rhodamine B or ATTO488 fluorescence was shown to dissociate from liposomes in human plasma [36]. In our study, we solved this problem by using very hydrophobic counterions to increase the hydrophobicity of the resulting dye salt and in this way improved encapsulation and strongly reduced leaking [37]. Moreover, high hydrophobicity of lumogen red also prevented leakage from PLGA in NPs, in line with previous data in serum and cells [26]. On the other hand, EET occurs only at high dye-counter ion concentrations [16]. Thus, the fluorescence of dye molecules leaking out of NPs dropped by several orders of magnitude and became undetectable. Therefore, unspecific fluorescence due to leaked dyes was not an issue in the current study.

A major challenge of the specific detection of NPs in the fixed brain is the strong auto-fluorescence of cerebral tissue after PFA exposure deriving, among others, from, mitochondria, collagen, or lipofuscin [38]. Currently autofluorescence quenching kits are reducing predominantly non-lipofuscin autofluorescence [39], however, cerebral autofluorescence is highly abundant of lipofuscin and is often particularly strong in structures of the same size and brightness as NPs. Therefore, broad emission spectrum cerebral auto-fluorescence often generates signals which very closely mimic NPs and may also generate false positive results. In the current study, we detected strong autofluorescence in neurons and endothelial cells, a finding well in line with the literature [40]. Subsequently, we avoided false positive identification of NPs by additional scanning of a channel outside the emission rage of NP fluorescence. Signals detected in both channels were regarded as autofluorescence and used to correct NP counts for false positive events.

Another point of discussion is whether single particles with a size of 70 nm, i.e. well below the resolution of a light microscope, i.e. approx. 250 nm, can be identified with the technology used in the current study. In fact, the size of an object observed with an optic system is determined by its physical boundaries, but also by its brightness because of the dispersion of light along the light path. This principle holds also true for very bright NPs like the ones created and used in the current study. Although their physical size is below the resolution of confocal or 2-photon microscopy, their brightness allowed us to most likely detect single particles in tissue sections and *in vivo*. Further experiments using novel technology of correlation electron and confocal fluorescence microscopy, however, will be needed to unambiguously address this issue.

Using ultra-bright NPs and autofluorescence correction, we demonstrate uptake of PF-68 coated PLGA NPs into the brain with high resolution and specificity. Our findings unravel, that the mechanism responsible for uptake of NPs into the brain is endocytosis by endothelial cells. Current study demonstrates that it is technically possible to track the transport of NPs on the cellular and subcellular level *in vivo* modulated by surface modifications. Therefore our findings pave the way for future studies on endothelial receptor-mediated transcytosis as well as other transport systems of NPs in cerebral vessels and may help to uncover the exact transport pathways of NPs reported to facilitate the transport of pharmacologically active molecules into the CNS [10, 12].

Coating of NPs with PF-68 was reported to reduce phagocytosis by liver and spleen macrophages [41] and we could confirm these results, though to a lower extent. More importantly, however, we demonstrated by *in vivo* confocal imaging at single-particle resolution that coating of NPs with PF-68 shifted the uptake of NPs from immune towards endothelial cells. This process occurred not only in the brain, but also in all other investigated organs, *i.e.* liver, spleen, kidney, and heart. Whether the undelaying mechanism is a longer plasma half-life or specific transport of the NPs is beyond the scope of the current investigation and needs to be addressed by follow-up studies. After all, these results also impressively demonstrate the potential of the currently developed experimental approach to specifically detect NPs in living tissue.

## 4. Conclusions

Taken together, we developed, highly fluorescent 70 nm PLGA NPs and demonstrate the bio-distribution of these drug delivery systems in the mouse *in vivo* by single- and multi-photon microscopy. Using this experimental approach, we demonstrate that uncoated PLGA NPs are mainly taken up by liver and spleen macrophages, while PF-68 coating shifts the uptake towards endothelial cells. In the brain, PF68-coated PLGA NPs end up in late endothelial endosomes. Thus, the combination of novel, highly fluorescent PLGA NP with high resolution *in vivo* imaging, and strategies to avoid unspecific autofluorescence allowed us to unambiguously detect the biodistribution of NPs down to the subcellular level with high accuracy, specificity, and resolution. This approach allowed us to identify PLGA nano-carrier coated by PF-68 able to target endothelial cells the brain. Further, our current technical approach *in vivo* may help to significantly close the existing translational gap between the preclinical and clinical evaluation of NPs.

## 5. Material and methods

### 5.1 Materials

Poly(D,L-lactide-co-glycolide) (PLGA, acid terminated, lactide:glycolide 50:50, M_w_ = 24,000-38,000), poloxamer 407 (Pluronic F-127) and poloxamer 188 (Pluronic F-68), propargylamine (98%), copper(II) sulfate pentahydrate (98.0%), sodium ascorbate (>98.0%), sodium azide (99%), sodium chloride (molecular biology grade), triethylamine (TEA, >99.5%), acetonitrile (anhydrous, 99.8%), dichloromethane (anhydrous, >99.8%), and N,N-dimethylformamide (absolute >99.8%) were purchased from Sigma-Aldrich. N,N-diisopropylethylamine (DIPEA, >99.0%), methanesulfonyl chloride (>99.7%), and 2-aminoethane sulfonic acid (taurine, >98.0%) were obtained from TCI. 1-hydroxybenzotriazole (HOBt, >99.0%), N-tetramethyl-O-(1H-benzotriazol-1-yl)uronium hexafluorophosphate (HBTU, 99.5%), and 1-[bis(dimethylamino)methylene]-1H-1,2,3-triazolo[4,5-b]pyridinium 3-oxid hexafluorophosphate (HATU, 99.8%) were purchased from chemPrep. Sodium phosphate monobasic (>99.0%, Sigma-Aldrich) and sodium phosphate dibasic dihydrate (>99.0%, Sigma-Aldrich) were used to prepare 20 mM phosphate buffer solutions at pH 7.4. MilliQ water was deionized using a Millipore purification system. R18/F5-TPB was synthesized from rhodamine B octadecyl ester perchlorate (Aldrich, >98.0%) and lithium tetrakis(pentafluorophenyl)borate ethyl etherate (AlfaAesar, 97%) through ion exchange followed by purification through column chromatography as described previously [16, 42]. N,N-Bis-(2,6-diisopropylphenyl)-1,6,7,12-tetraphenoxy-3,4,9,10-perylenebis(dicarboximide) (Lumogen Red) was purchased from ORGANICA® Feinchemie GmbH Wolfen.

#### 5.1.1 Sulfonated lissamine-alkyne

was synthesized as described previously [29, 43]. Briefly, lissamine, (100 mg, 0.16 mmol, 1 eq), propargylamine (11 mg, 0.19 mmol, 1.2 eq), HATU (76 mg, 0.16 mmol, 1 eq), and DIPEA (127 μL, 0.75 mmol, 5 eq) were solubilized in anhydrous DMF (5 mL) under argon. After stirring at room temperature for 24h, the mixture was dried under reduced pressure at 65°C and then diluted with DCM (20 mL) and extracted four times with water. The combined organic phases were dried over sodium sulfate and concentrated *in vacuo* and, subsequently, the residue was purified by flash chromatography eluting with DCM/MeOH (99:1) to give 89.5 mg of a pink solid (yield: 54%).

^1^H NMR (400 MHz, MeOD): δ = 8.65 (1H, s, ar. CH) + 8.03 (1H, d, ar. CH) + 7.29 (1H, d, ar. CH) + 7.16 (2H, d, ar. CH) + 6.85 (2H, d, ar. CH) + 6.73 (2H, s, ar. CH) ~9H, 3.95 (2H, s, -NCH_2_C-), 3.51-3.65 (8H, m, -NC*H_2_*CH_3_), 2.45 (2H, t, -NCOCH_2_-), 2.27 (1H, s, CCH), 1.23-1.34 (12H, m, -NCH_2_C*H_3_*).

#### 5.1.2 Dimesyl derivative of PF-127

PF-127 (6.3 g, 0.5 mmol, 1 eq) was solubilized in DCM (25 mL) and cooled to 0°C. Next, TEA (420 μL, 3 mmol, 6 eq), and methanesulfonyl chloride (234 μL, 3 mmol, 6 eq) were added. The reaction mixture was kept under stirring at 0°C for 3 h and then at room temperature overnight. The solution was dried under reduced pressure at 40°C for 30 minutes. The obtained solid was redispersed in water and purification is carried out by means of dialysis against water (48 h) to give 5.2 g of a white solid (yield: 82%).

^1^H NMR (400 MHz, MeOD): δ = 3.81-3.78 (4H, m, -SOCH_2_*CH_2_*-), 3.77-3.61 (-O*CH_2_CH_2_*O-) + 3.59-3.51 (m, -O*CH_2_*CH-) + 3.43-3.38 (m, -*CH*CH_3_) ~1000H, 3.14 (6H, s, C*H_3_*SOO^−^), 1.16 (~195H, m, -CH*CH_3_*).

#### 5.1.3 Diazide derivative of PF-127

The dimesyl derivative of PF-127 (5.2 g, 0.41 mmol, 1 eq) and sodium azide (165 mg, 2.46 mmol, 6 eq) were solubilized in acetonitrile (25 mL) and heated under reflux for 48 h. The obtained solid was redispersed in water and purification was carried out by means of dialysis against water (48h) to give 4.6 g of a white solid (yield: 87%).

^1^H NMR (400 MHz, MeOD): δ = 3.77-3.61 (-O*CH_2_CH_2_*O-) + 3.59-3.51 (m, -O*CH_2_*CH-) + 3.43-3.38 (m, -*CH*CH_3 +_ *CH_2_*N_3_) ~1000H, 1.16 (~195H, m, -CH*CH_3_*).

### 5.2 Click reaction of lissamine on pluronic

Sodium ascorbate (13 mg, 0.074 mmol, 16.5 eq in 100 μL of water) was added to a Copper(II) sulfate pentahydrate (10 mg, 0.04 mmol, 9 eq in 100 μL of water). Then the solution was added in a mixture of diazide derivative of PF-127 (55 mg, 0.0045 mmol, 1 eq), and sulfonated lissamine-alkyne (9 mg, 0.013 mmol, 2.9 eq) dissolved in anhydrous DMF (5 mL). The heterogeneous mixture was stirred vigorously for 24 hours at 55°C under argon. The reaction mixture was dried under reduced pressure at 60°C, diluted in DCM, and then extracted four times with water. The combined organic phases were dried over sodium sulfate and purified by size exclusion chromatography to the final purified PF-127 conjugate with corresponding dye.

Sulfonated lissamine-alkyne PF-127 conjugate: 31 mg of a pink solid (yield: 57%), ^1^H NMR (400 MHz, MeOD): δ = 8.65-6.70 (~11H, m, ar CH), 3.77-3.61 (-OCH_2_CH_2_O-) + 3.59-3.51 (m, -OCH_2_CH-) + 3.43-3.38 (m, -CHCH_3_) ~1000H, 1.04 (~195H, m, -CHCH_3_). Degree of modification 61% (percentage of sulfonate lissamine-alkyne linked to pluronic).

### 5.3 Nanoparticle preparation

Stock solutions of the PLGA in acetonitrile were prepared at a concentration of 10 mg·mL^−1^ and, then, diluted with acetonitrile to 4 mg·mL^−1^. The desired amount of R18/F5-TPB (typically 1 wt% relative to the polymer) or lumogen red (1wt%) was added. Under constant shaking, 50 μL of the polymer solution were quickly added using a micropipette (Thermomixer comfort, Eppendorf, 1100 rpm, 21°C) to 450 μL of milliQ water containing a chosen concentration of sodium chloride. Particle size was tuned by modulating sodium chloride concentration from 0 to 50 mM. The particle suspension was then quickly diluted 5-fold with water. For stabilization of NPs, different amounts of 1 or 0.1 mg mL^−1^ solutions of PF-127 or PF-68 were added under stirring to the NP solutions.

#### 5.3.1 Instrumentation

The size and polydispersity index (PDI) measurements of the NPs were performed by DLS on a Zetasizer Nano series DTS 1060 (Malvern Instruments S.A.), using a 633 nm laser, which excludes any light excitation of our dye-loaded NPs. Absorption and emission spectra were recorded on a Cary 400 Scan ultraviolet-visible spectrophotometer (Varian) and a FluoroMax-4 spectro-fluorometer (Horiba Jobin Yvon) equipped with a thermostated cell compartment, respectively. For standard recording of fluorescence spectra, the excitation wavelength was set at 530 nm and emission was recorded from 540 to 800 nm. The fluorescence spectra were corrected for detector response and lamp fluctuations.

Fluorescence correlation spectroscopy (FCS) measurements were carried out on a two-photon platform based on Olympus IX70 inverted microscope. Two-photon excitation at 780 nm (5 mW laser output power) was provided using a mode-locked Tsunami Ti: sapphire laser pumped using a Millenia V solid state laser (Spectra Physics). The measurements were performed in an eight-well Lab-Tek II coverglass system, using 300 μL volume per well. The focal spot was set about 20 μm above the coverslip. The normalized autocorrelation function, G(τ), was calculated online using an ALV-5000E correlator (ALV, Germany) from the fluorescence fluctuations, δF(t), by G(τ)= =<δF(t)δF(t +τ)>/<F(t)>^2^ where t is the mean fluorescence signal and τ is the lag time. Assuming that NPs diffuse freely in a Gaussian excitation volume, the correlation function, G(τ), calculated from the fluorescence fluctuations was fitted according to Thompson [44]:

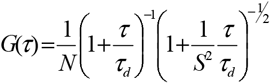

where τ_d_ is the diffusion time, N is the mean number of fluorescent species within the two-photon excitation volume, and S is the ratio between the axial and lateral radii of the excitation volume. The excitation volume is about 0.34 fL and S is about 3 to 4. The typical data recording time was 5 min. Measurements were done with respect to a reference 5(6)-carboxytetramethylrhodamine (TMR) (Sigma-Aldrich) in water.

#### 5.3.2 Single-particle measurements on surfaces

NPs were immobilized on glass surface of LabTek slides as previously described [16]. Single-particle measurements were performed in the epi-fluorescence mode using Nikon Ti-E inverted microscope with CFI Plan Apo 60x oil (NA = 1.4) objective. The excitation was provided by light emitting diodes (SpectraX, Lumencor) at 550 nm at power density of 1.3 W·cm^−2^. The fluorescence signal was recorded with a Hamamatsu Orca Flash 4 camera. Integration time for a single frame was 200 ms. The single-particle analysis was performed using Fiji software, similarly to the previously described protocol [27]. Briefly, particle locations were detected through a FIJI routine applied to a projection (maximum intensity) of 10 frames using an appropriate threshold. The mean intensities of circular regions of interest with a diameter of 6 pixels around the found particle locations were then measured. Background subtraction was then achieved by measuring the mean intensities in circular bands around the circular regions of interest and subtracting them. Finally, integrated intensity in all circular regions of interest (for the first 5 frames) was calculated and used for building the histograms of particle intensities.

### 5.4 Kinetics of NPs in the brain vasculature

All animal experiments were conducted in accordance with institutional guidelines and approved by the Government of Upper Bavaria. For real-time *in vivo* multiphoton imaging we used an upright Zeiss LSM710 microscope as described [29]. Briefly, 8-week old C56/Bl6N mice were anesthetized intraperitoneally with a combination of medetomidine (0.5 mg·kg^−1^), fentanyl (0.05 mg·kg^−1^), and midazolam (5mg kg^−1^) (MMF). Then animals were endotracheally intubated and ventilated in a volume controlled mode (MiniVent 845, Hugo Sachs Elektronik, March-Hungstetten, Germany) with continuous recording of end◻tidal pCO2. The body temperature was monitored throughout the experiment and maintained by a rectal probe attached to a feedback◻controlled heating pad. Mouse femoral artery was catheterized in order to measure mean arteriolar blood pressure and inject solutions directly into the blood system. The intravital imaging was performed through a rectangular 4×4◻mm glass cranial window at right fronto-parietal cortex, located 1◻mm lateral to the sagittal suture and 1◻mm frontal to the coronal suture made as described [29]. Afterwards, mice were placed on the multiphoton microscope adapted for intravital imaging of small animals. During imaging, first, mice were injected with FITC◻dextran 3μL·g^−1^ to locate the region of interest (ROI). Upon identifying the ROI and scanning the baseline image, animals were administered with 7.5 μL·g^−1^ of PLGA NPs: bare (group of 3 animals) or F68 coated (group of 6 animals). The scanning was performed at 150 μm depth with laser power 4%-20%, laser wavelength 800 nm and image acquisition was done through GAASP detector with LP◻570 nm filter and master gain 600 for the FITC channel and LP◻570 nm for the NPs channel with master gain 530.

#### 5.4.1 Imaging analysis

The fluorescence analysis was performed using FIJI software. The Z-stack was converted to maximum intensity projection and the channels were split. The integrated density value of the each timepoint was divided (normalized) on the integrated density value of the baseline. To prevent fluctuation of the signal between animals, the signal from NPs channel was also divided (normalized) on FITC channel. The Two-way ANOVA statistical analysis with Tukey correction for multiple comparison was performed using Graph Pad Prism 8.0.2. The video data as well as 3D-reconstraction was prepared using Imaris® software.

### 5.5 Evaluation of NPs bio-distribution in fixed tissue

C57Bl/6N male mice (23-25g) were injected intravenously (IV) with 7.5 μl·g^−1^ of PBS, PLGA-bare or PF-68-coated PLGA NPs solutions (*n*=3). 60 min post-injection, animals were anesthetized with a triple combination MMF (i.p.; 1 ml per 100 g body mass for mice). Upon the absence of the paw reflexes, the abdomen and chest were open and 25 G needle was inserted into the left ventricle for the perfusion with 0,9 % NaCl followed by 4 % PFA. The brain, liver, spleen, heart and kidney were extracted and fixed overnight in 4 % PFA.

#### 5.5.1 Mouse tissue processing and staining

##### 5.5.1.1 Brains

The sectioning of brains into 50μm slices was done by Leica Vibratome, which were subsequently collected and kept in 0.1 m phosphate-buffered saline (PBS). The frontal, middle and caudal parts of each brain were taken for the staining. The free-floating sections were permeabilized in 24-well plates with 0.1% Tween20© (Carl Roth GmbH + Co. KG, Karlsruhe, Germany) in 0.1 m PBS and blocked with 10 % goat serum in 0.1 m PBS. Staining was performed with 1:1500 anti-Laminin (Rabbit, Sigma, #L9393) and anti-rabbit coupled to Alexa-fluor 647 (donkey anti-rabbit, Jackson Immuno Research, # 711-606-152, 1.5mg·mL^−1^ in 50% glycerol) as well as 1:200 anti-LAMP1 (rat, Santa Cruz, #sc-19992) and anti-rat coupled to Alexa-fluor 647 (donkey anti-rat, Jackson Immuno Research, #712-606-150, 1.5mg·mL^−1^ in 50% glycerol). Nuclei were stained with 4′,6-Diamidin-2-phenylindol (DAPI, Invitrogen, #D1306) 1:5000. DAPI, both primary and secondary antibodies were applied in 0.1 M PBS, because long-time exposure with triton or tween led to substantial leakage of NPs out of the tissue. Sections were mounted on microscope slides (Menzel-Gläser Superfrost® Plus, Thermo Fisher Scientific, #3502076) and covered with a coverslip (Menzel-Gläser 24–60 mm, #1, BB024060A1, Wagner und Munz) using aqueous mounting medium (CC/MountTM, Sigma-Aldrich, #C9368-30ML).

##### 5.5.1.2 Internal organs

Free floating 50 μm sections of brain, liver, kidney, spleen and heart were cut vibratome. Then sections were permeabilized for 30 minutes in PBS Tween 20, blocked for 60 minutes in 10% goat serum in PBS and then stained with the primary antibody for 12 hours at 4°C. The following primary antibodies were used: iba-1 (rabbit, Wako, #019-19741, 1:100), collagen IV (rabbit, Abcam, #ab19808, 1:100). After incubation sections were washed in PBS and incubated with the following secondary antibodies: anti-rabbit coupled to Alexa-fluor 594 (goat anti-rabbit, Thermo Fisher Scientific, #A-11012). Nuclei were stained with 4’,6-Diamidin-2-phenylindol (DAPI, Invitrogen, #D1306) 1:10,000 in 0.01 M PBS.

#### 5.5.2 Imaging and analysis of fixed tissue

##### 5.5.2.1 Brains

The whole hemisphere images were acquired using a Zeiss confocal microscope with 40x magnification (objective: EC Plan-Neofluar 40x/1.30 Oil DIC M27) with an image matrix of 512×512 pixel, a pixel scaling of 0.593×0.593 μm and a depth of 8 bit. Confocal-images were collected in tile scan Z-stacks 10 slices with a slice-distance of 2 μm. Single particles images were acquired with 100x magnification (alpha Plan-Apochromat 100x/1.46 Oil DIC M27 Elyra) with an image matrix 2048×2048, a pixel scaling of 0.059 μm x×0.059 μm and a depth of 8 bit. Morphological analysis was performed using FIJI software. The whole brain hemisphere was selected as a region of interest and the channels were split. The mean grey value of the rhodamine channel was divided (normalized) on the mean grey value of the DAPI channel. The One-way ANOVA statistical analysis with Tukey correction for multiple comparison was performed using Graph Pad Prism 8.0.2. Analysis of single particles in the brain parenchyma was performed using green 488 channel as a reference for the auto-fluorescence. Only red dotty structures that were strongly not co-localized with the green 488 channel had been considered as NPs. On merged image, the co-localized dots had yellowish colour, while NPs were red. The quantification of the autofluorescent particles (*n*=3, ROI=5): the number of co-localized (yellow colour) foci in the ROI was divided on the ROI’s area and cell numbers. The quantification of the single particles (*n*=3, ROI=5): the number of non-co-localized (red colour) foci in the ROI was normalized to the ROI area and cell numbers. The unpaired t-test was used for statistical analysis using Graph Pad Prism 8.0.2. The video data as well as 3D-reconstraction was prepared using Imaris® software.

##### 5.5.2.2 Internal organs

Imaging was performed using confocal microscopy (ZEISS LSM 900, Carl Zeiss Microscopy GmbH, Jena Germany). For colocalization analysis, 40x magnification (objective: EC Plan-Neofluar 40x/1.30 Oil DIC M27) was used with an image matrix of 512×512, a pixel scaling of 0.415×0.415 μm and a depth of 16 bit. Three to four ROIs per animal were collected in z-stacks as tile scans with a slice-distance of 1.07 μm and a total range of 5.35 μm.

For the co-localization analysis, we exploited, ImageJ compatible, JACoP Plugin [45]. Z-stacks were imported into the software in split into individual channels. The rhodamine channel for nanoparticles and the far red channel for either iba-1 or collagen IV were selected for co-localization analysis and thresholds were set using Costes’ automatic threshold as described previously [46]. Finally, the plugin calculates colocalization using the Manders’ overlap coefficient, established by Manders *et al.* [34]. This coefficient represents the ratio of summed intensities of particles in the predefined green and red channels and therefore indicates the proportion of NPs overlapping with iba-1 or collagen IV over their whole intensity. The Manders coefficient varies between 0 and 1, with 0 meaning no overlap and 1 being perfect colocalization. This can be done throughout the Z-stack without having to take the maximum intensity projection, therefore reducing signal loss through overlay of NPs.

## Supporting information

Supplements contain a Table,2 Figures and description of movies

## Supporting information

Supporting Information containing figures and movies are available at website.

## Acknowledgements

This project was funded by the Alexander von Humboldt Foundation, the European Union Horizon 2020 research and innovation program under the Marie Skłodowska-Curie grant agreement No 794094, the European Research Council ERC Consolidator grant BrightSens 648528, the Agence National de Recherche JC/JC grant “Supertrack” ANR-16-CE09-0007, the DFG under the Munich Cluster of Systems Neurology (Synergy), and ERA-NET Neuron TRAINS.

