## Supplements contain a Table,2 Figures and description of movies for "Dynamic tracing using ultra-bright labelling and multi-photon microscopy identifies endothelial uptake of poloxamer 188 coated poly(lactic-co-glycolic acid) nano-carriers *in vivo*"

**Cerebral biodistribution and real-time particle tracking *in vivo* of ultrabright fluorescent PLGA nano-carriers**

**Table S1.** Polydispersity indicis (PDI) of PLGA NPs with different fluorophores and coatings.

| **PDI** | **Bare** | **PF-127 (3000nM)** | **PF-68 (3000 nm)** |
| --- | --- | --- | --- |
| **Lumogen Red 1%** | 0.09 ±0.09 | 0.072±0.013 | 0.09±0.02 |
| **R18/F5-TPB 1%** | 0.09 ±0.01 | 0.08±0.02 | 0.10±0.02 |

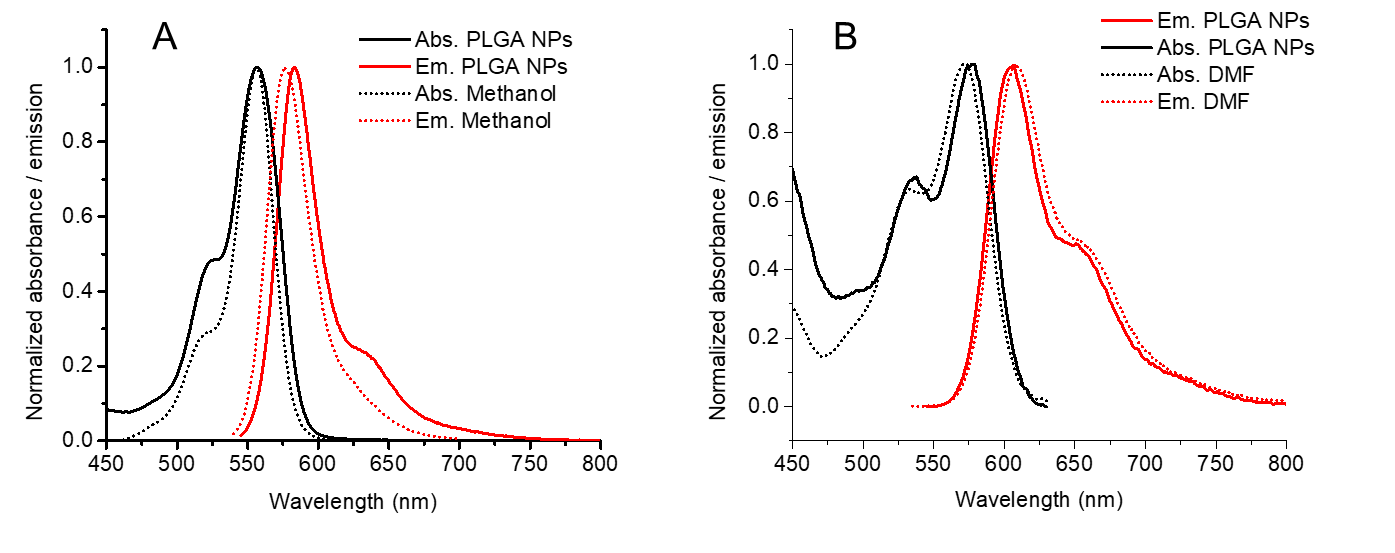

**Figure S1.** Normalized absorption (black) and emission (red) spectra of R18/F5-TPB (A) and Lumogen Red (B) in PLGA NPs (solid lines) and in an organic solvent (dotted lines).

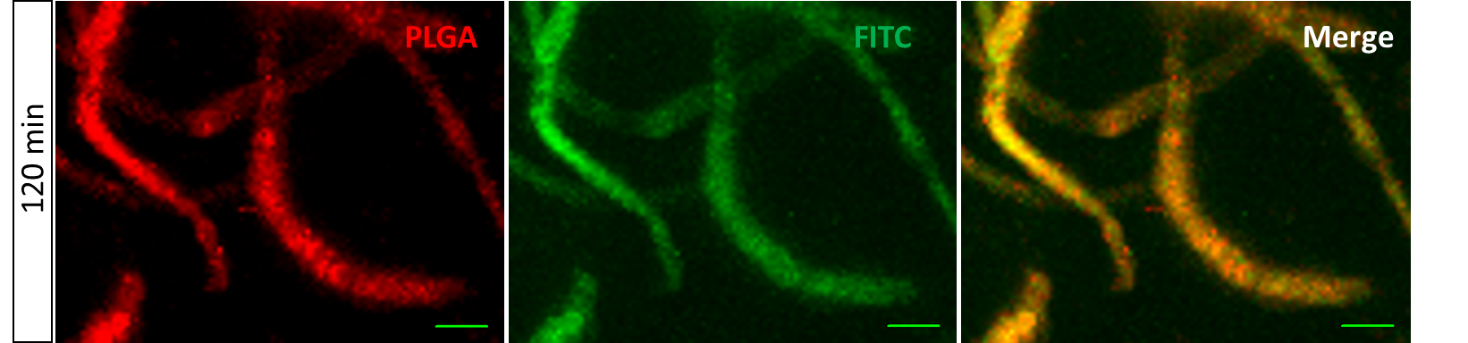

**Figure S2.** Accumulation of PLGA NPs in the vessels’s wall 120 minutes post-injection. Scanned images 120 minutes after injection of FITC-dextran 2000kDa and coated PLGA-F68 NPs. Z-stack, maximum intensity projection. Scale bar - 10 μm.

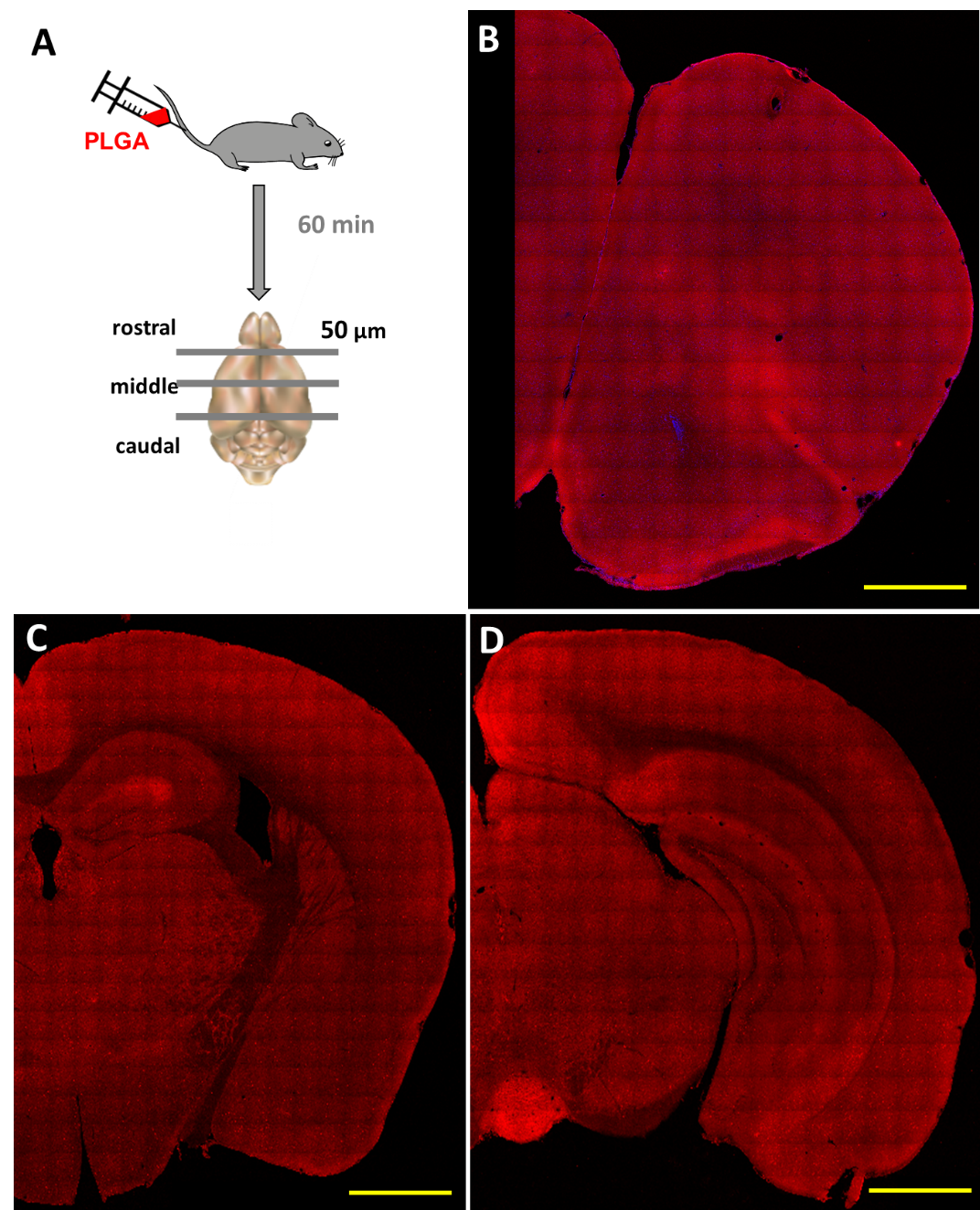

**Figure S3.** Fixed brain analysis. **A:** Experimental design. **B-D**: Representative 40x confocal images of whole brain coronal sections (rostral (B); middle (C); caudal (D)) of mouse injected with coated nanoparticles. Maximum intensity projection of rhodamine channel of tile scans. Scale bar - 1 mm.

Supporting movies

**Movie S1**. Bare PLGA particles circulating in the mouse brain vasculature. Scanned images 5 min after injection of NPs solution. Rendered Z-stack, 150 μm depth. Laser power 3.5%-10%. Green - FITC-dextran (2000kDa), red foci – bare PLGA NPs, 70 nm. Scale bar - 50 μm.

**Movie S2**. PF-68 costed PLGA particles circulating in the mouse brain vasculature. Scanned images 5 min after injection of NPs solution. Rendered Z-stack, 150 μm depth. Laser power 3.5%-10%. Green - FITC-dextran (2000kDa), red – coated PLGA NPs, 70 nm. Scale bar - 50 μm.

**Movie S3.** 3D reconstruction of brain capillary with accumulated PF-68 coated PLGA NPs. Rhodamine channel shows non-specific red foci. Auto-fluorescent channel shows auto-fluorescent foci of the brain parenchyma. Red arrow indicates the PLGA NPs, while white arrow - auto-fluorescence as a co-localization between rhodamine and auto-fluorescent channels. Laminin stains basal membrane, which outlines the spatial distribution of NPs.

**Movie S4.** 3D reconstruction of LAMP1 positive cell compartments; red – NPs; green – autofluorescence; white and red – a co-localization of NPs and LAMP1; white and green – a co-localization of autofluorescent focus and LAMP1.

**Movie S5.** 3D reconstruction of PLGA NPs uptaken by Kuppfer cell (Iba-1 positive) of the mouse injected by bare NPs.

**Movie S6.** 3D reconstruction of PLGA NPs uptaken by Kuppfer cell (Iba-1 positive) of the mouse injected by PF-68 coated NPs.
